# Unraveling the genetic architecture of the adaptive potential of *Arabidopsis thaliana* to face the bacterial pathogen *Pseudomonas syringae* in the context of global change

**DOI:** 10.1101/2022.08.26.505380

**Authors:** Claudia Bartoli, Mylène Rigal, Baptiste Mayjonade, Fabrice Roux

## Abstract

Phytopathogens are a continuous threat for global food production and security. Emergence or re-emergence of plant pathogens is highly dependent on the environmental conditions affecting pathogen spread and survival. Under climate change, a geographic expansion of pathogen distribution poleward has been observed, potentially resulting in disease outbreaks on crops and wild plants. Therefore, estimating the adaptive potential of plants to novel epidemics and describing its underlying genetic architecture, is a primary need to propose agricultural management strategies reducing pathogen outbreaks and to breed novel plant cultivars adapted to pathogens that might spread in novel habitats under climate change. To address this challenge, we inoculated *Pseudomonas syringae* strains isolated from *Arabidopsis thaliana* populations located in south-west of France on the highly genetically polymorphic TOU-A *A. thaliana* population located east-central France. While no adaptive potential was identified in response to most *P. syringae* strains, the TOU-A population displays a variable disease response to the *P. syringae* strain JACO-CL belonging to the phylogroup 7 (PG7). This strain carried a reduced T3SS characteristic of the PG7 as well as flexible genomic traits and potential novel effectors. GWA mapping on 192 TOU-A accessions inoculated with JACO-CL revealed a polygenic architecture. The main QTL region encompasses two *R* genes and the *AT5G18310* gene encoding for ubiquitin hydrolase, a target of the AvrRpt2 *P. syringae* effector. Altogether, our results pave the way for a better understanding of the genetic and molecular basis of the adaptive potential in an ecologically relevant *A. thaliana* – *P. syringae* pathosystem.

## INTRODUCTION

Pathogens are a threat both for crops and for wild species. Yield losses resulting from pathogen attacks can be up to several tens of percent in crops (Oerke, 2006; McDonald and Stukenbrock, 2016; Savary et al., 2019), thereby threatening global food security (Ristaino et al., 2021; Savary et al., 2019). Experimental evidences indicate that naturally infected wild species plants can harbor a significant decrease in the number of offspring (Jarosz and Davelos, 1995; Traw et al., 2007; Frachon & Roux 2022). A major challenge in plant breeding and ecological genomics is to identify the genetic and molecular bases of natural variation in disease resistance (Bartoli and Roux, 2017; Roux and Bergelson, 2016). The characterization of the genetic architecture of disease resistance might have enormous practical implications by increasing crop yield and quality (Deng et al., 2020; Karasov et al., 2020; Li et al., 2020), and lead to fundamental insights in the prediction of evolutionary trajectories of natural populations (Karasov et al., 2014; Roux et al., 2014). This is particularly relevant in the current climate change that is likely to favor conditions for pathogen spread and their adaptation potential in comparison with sessile plants (Bebber et al., 2014; Bergot et al., 2004; Garrett et al., 2006; Pautasso et al., 2012; Tylianakis et al., 2008). For instance, temperature elevation is expected to favor the emergence of new pathogens and to increase the severity and occurrence of epidemics (Bebber, 2015; Desaint et al., 2021; Elad and Pertot, 2014; McDonald and Stukenbrock, 2016). In addition, climate change scenarios predict an increase in epidemic severity and a northwards geographic expansion of pathogen distribution in Western Europe (Bergot et al., 2004; Evans et al., 2008; Prank et al., 2019). Accordingly, a global warming-driven movement poleward has already been observed for a large number of pathogens (Bebber et al., 2013).

*Pseudomonas syringe* is a ubiquitous phytopathogenic bacterium belonging to a species complex including strains with a large host range and adaptability to a wide spectrum of environment outside of crop fields. Non-agricultural strains of *P. syringae* have been isolated from wild plant species (including Arabidopsis), snow, rain, rivers, litter etc. (Monteil et al., 2014b; Morris et al., 2013), corroborating the great adaptability of strains of the *P. syringae* complex to colonize a large range of environments. Additionally, air masses and precipitations are a way for *P. syringae* translocation over large geographic distances (Monteil et al., 2014a). *P. syringae* strains growing in icy and hostile conditions similar to cloud environment, were also shown to acquire exogenous DNA faster than other bacterial species. This suggests that *P. syringae* strains are able to improve their genomic content and thus their adaptability while moving in the air masses (Blanchard et al., 2017). Under these circumstances, it is not surprising that *P. syringae* outbreaks originate from hosts and habitats different than the final host where the disease is reported (Koike et al., 2017). The wide host range coupled with the great adaptability of *P. syringae* and its ability to colonize an incredibly diverse range of habitats, question the impact that the climatic change can play on the dispersal of this phytopathogen. Therefore, understanding the adaptability of *P. syringae* strains isolated from the south to colonize and infect plant populations adapted to colder conditions is a pertinent question to answer to better understand and control disease outbreaks.

In this study, we aimed at estimating the adaptive potential, and describing the related underlying genetic architecture, of the annual wild species *Arabidopsis thaliana* to face attacks from more southern strains of the *P. syringae* complex. Besides being a model in plant genetics and molecular biology, *A. thaliana* is above all a wild species inhabiting contrasting environments for diverse abiotic (e.g. climate, soil physico-chemical properties) and biotic (e.g. microbial communities, plant communities) factors (Bartoli et al., 2018; Brachi et al., 2013; Frachon et al., 2018; Frachon et al., 2019; Thiergart et al., 2020). In addition, disease incidence has been commonly observed in natural populations of *A. thaliana* (Barrett et al., 2009; Roux and Bergelson, 2016). For instance, a survey of 163 natural populations in south-west of France reported that 72.7% of plants presented disease symptoms, with each of the two most abundant bacterial pathogens (namely the *Pseudomonas syringae* complex and *Xanthomonas campestris*) being detected in more than 90% of the populations by a metabarcoding approach (Bartoli et al., 2018).

More precisely, we aimed at testing the adaptive genetic potential of the TOU-A local mapping population located in Burgundy (east of France) to face the individual attack of eight *P. syringae* strains isolated from natural populations of *A. thaliana* located ∼400km south-west of the TOU-A population. We chose to use this local mapping population for its several advantages. First, located under a ∼350 meter long electric fence, the TOU-A population is highly polymorphic (Frachon et al., 2017). Based on the whole-genome sequences of 195 TOU-A accessions, more than 1.9 million single nucleotide polymorphisms (SNPs) were detected, only 5.6 times less than observed in a panel of 1,135 worldwide accessions (Frachon et al., 2017). Secondly, extensive genetic variation for resistance to diverse pathogens was detected when conducting experiments in controlled laboratory conditions (Debieu et al., 2016; Frachon et al., 2017; Huard-Chauveau et al., 2013) or directly in the native habitat of the TOU-A population (Roux & Frachon 2022). Thirdly, a short linkage disequilibrium (LD) decay, below 3 kb, combined with a strongly reduced confounding effect of population structure allows fine-mapping of genomic regions associated with phenotypic variation (Baron et al., 2015; Brachi et al., 2013; Frachon et al., 2017; Libourel et al., 2021), which in turn leads to the cloning and functional validation of QTLs of disease resistance (Aoun et al., 2020) .

## RESULTS

### The adaptive genetic potential of the TOU-A population to face attacks from more southern *P. syringae* strains is limited

To estimate the adaptive genetic potential of the TOU-A population to face attacks from more southern *P. syringae* strains, we first challenged seven accessions covering the genomic space of the TOU-A population (Frachon et al., 2017) with seven *P. syringae* strains isolated from contrasting habitats from south-west of France, i.e. 0105-Psy-JACO-CL, 0106-Psy-RADE-AL, 0111-Psy-RAYR-BL, 0114-Psy-NAUV-BL, 0117-Psy-NAZA-AL, 0124-Psy-SAUB-AL and 0132-Psy-BAZI-AL hereafter named BAZI-AL, JACO-CL, NAZA-AL, NAUV-BL, RADE-AL, RAYR-BL and SAUB-AL (Figure 1a). In our phenotypic assay, we also included the reference *A. thaliana* accession Col-0. By aligning a fragment of the *citrate synthase* (*cts*) housekeeping gene of the seven strains with reference sequences (Berge et al., 2014), a phylogenetic inference shows that four strains (JACO-CL, NAUV-BL, RADE-AL, and SAUB-AL) belong to the *P. syringae* Phylogroup (PG) 7 *clade a* (Figure 1b) and are closely related to the TA043 strain isolated from *Primula officinalis* (Morris et al., 2010). In the past, PG7 was known under the name of *P. viridiflava* (Bartoli et al., 2014; Parkinson et al., 2011), a phytopathogen with pectolytic activity and described to cause disease on a wide-range of crops (Conn, 1993; González et al., 2003; Goumans and Chatzaki, 1998; Morris, 1992). Re-classification of *P. viridiflava* demonstrated that this phytopathogen is defined by two *P. syringae* PGs (7 and 8), PG7 *clade a* being the most abundant in both agricultural and non-agricultural habitats (Lipps et al., 2022; Lipps and Samac, 2022). Several herbaceous plants, including *A. thaliana*, have been described as natural hosts of PG7. Accordingly, PG7 *clade a* is one of the most abundant bacterial OTU inhabiting *A. thaliana* leaves (Bartoli et al., 2018; Goss, 2004; Jackson et al., 1999). The three other strains belong to PGs 2, 4 and 13 of *P. syringae sensu stricto* (BAZI-AL, NAZA-AL and RAYR-BL) (Figure 1b). We also included as a control the strain SIMO-AL, a Pseudomonas strain that is phylogenetically distant from the *P. syringae* complex.

**Figure 1.**
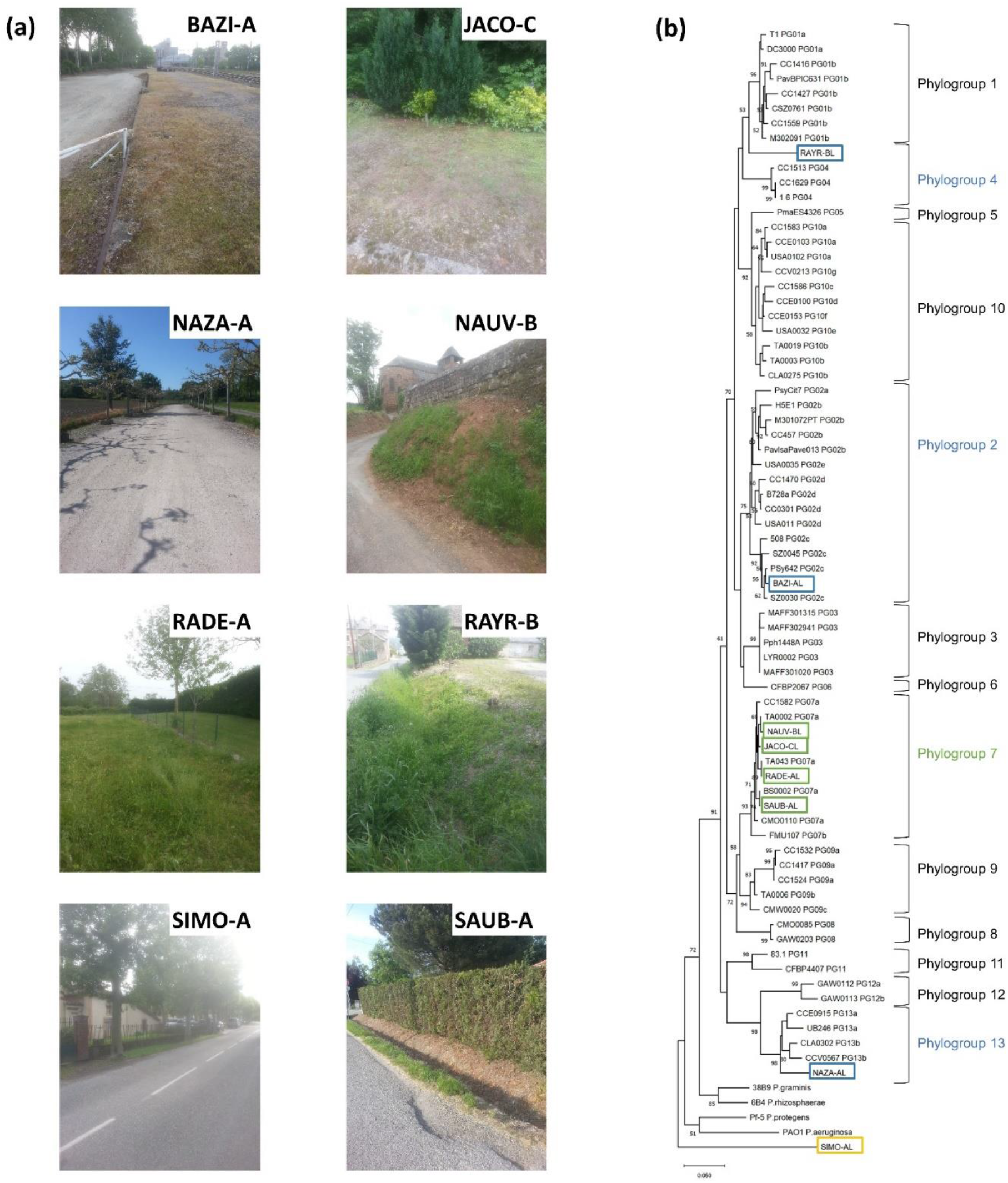
Ecological habitats and phylogenetic affiliation of the eight natural strains used in this study. **(a)** Diversity of habitats from which the eight Pseudomonas strains have been isolated in south-west of France. BAZI-A: Baziège, JACO-C: Jacoy (Boussenac), NAZA-A: Saint-Pierre-de-Najac, NAUV-B : Nauviale, RADE-A : Sainte-Radegonde, RAYR-B : Rayret (Cassagnes-Bégonhès), SIMO-A : Simorre, SAUB-A : Saubens. **(b)** Phylogenetic tree (Neighbor Joining, NJ) based on the *cts* sequences (406 bp) of the 64 strains proposed by Berge *et al*., (2014) for the rapid identification and classification of the seven strains belonging to the *Pseudomonas syringae* complex.

When tested on eight local accessions of *A. thaliana* from south-west of France, very few or no symptoms were observed in response to BAZI-AL, JACO-CL, NAZA-AL and SIMO-AL, whereas strong symptoms were observed in response to the RAYR-BL strain (Bartoli et al., 2018). Milder symptoms were observed in response to the three *P. viridiflava* strains NAUV-BL, RADE-AL and SAUB-BL (Bartoli et al., 2018).

Similar results were observed with the seven TOU-A accessions (with or without considering Col-0) 1, 2, 3 and 4 days after inoculation (dai), with the exception of the *P. viridiflava* JACO-CL strain that was more aggressive on the TOU-A accessions than on the accessions from south-west of France (Figure 2, Table S1, Supplementary Data Set 1). The absence of aggressiveness of the BAZI-AL strain is congruent with the absence of a canonical and complete T3SS in PG2 *clade c* strains (Dillon et al., 2019). In fact, *P. syringae clade 2c* is dominated by non-pathogenic *P. syringae* strains with an atypical T3SS similarly organized to the T3SS of S-PAI (Single-Pathogenicity Island) PG7 strains (Berge et al., 2014). This indicates that PG *clade 2c* strains probably tent toward low virulence and epiphytic interactions (Hirano and Upper, 2000). Likewise, the absence of symptoms observed when inoculating the NAZA-AL belonging to the PG13 is coherent with this phylogroup including mostly non-pathogenic strains from non-plant substrates with an atypical and reduced T3SS more related to the S-PAI found in the PG7 strains (Berge et al., 2014).

**Figure 2.**
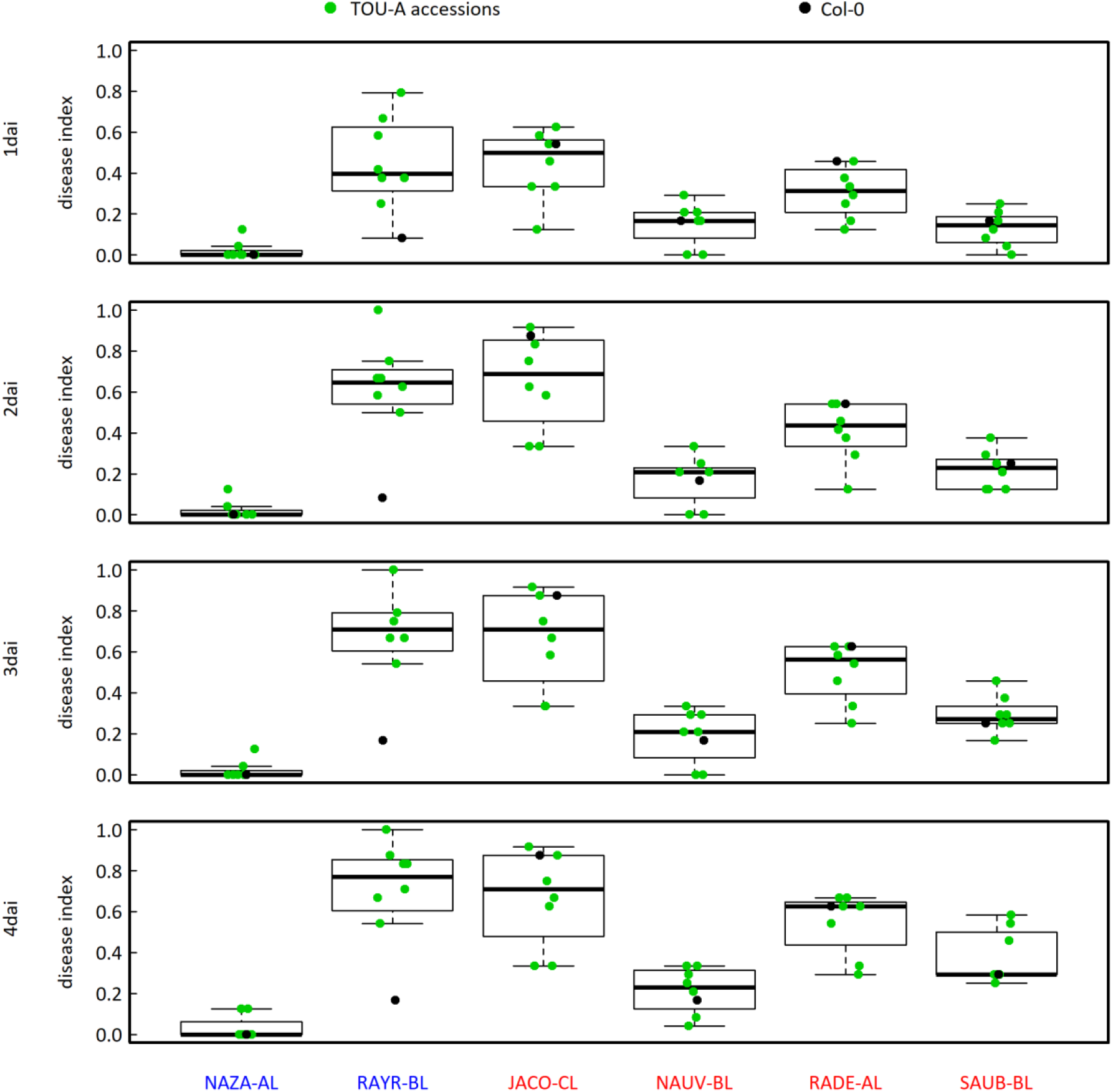
Genetic variation of virulence between six belonging to the *Pseudomonas syringae* complex, across eight accessions of *A. thaliana* at 1dai, 2 dai, 3 dai and 4 dai. The four strains of *P. viridiflava* are in red while two strains of *P. syringae* are in blue. No symptoms was observed on any plant for the two other strains of *P. syringae* (BAZI-AL and SIMO-AL, Supplementary Table S1). Dots corresponds to the genotypic values of the seven TOU-A accessions (green) and the reference accession Col-0 (black).

On the other hand, the high virulence of the RAYR-B strain on diverse sets of *A. thaliana* accessions is likely mediated by the T3E HopBJ1, previously identified and functionally characterized in this strain as inducing ROS accumulation and cell death both in Arabidopsis and in tobacco (Fuenzalida-Valdivia et al., 2022; Zavala et al., 2022).

Importantly, no genetic variation was detected between the seven TOU-A accessions in response to the three *P. viridiflava* strains NAUV-BL, RADE-AL and SAUB-BL, which can induce severe symptoms (Figure 2, Table S2). This result suggests a limited genetic potential of the TOU-A population to face attacks from a non-negligible fraction of more southern strains of the *P. syringae* complex. However, we caution that the restricted number of TOU-A accessions used in this experiment may not reflect the entire range of genetic variation present within the TOU-A population.

On the other hand, significant genetic variation was detected between the TOU-A accessions in response to the strains JACO-CL and RAYR-BL (Figure 2, Table S2). This is in line with some studies (mainly in trees) reporting genetic variation in host plant resistance to emerging pathogens (Ahrens et al., 2019; Alcaide et al., 2020; Elvira-Recuenco et al., 2014; Kjær et al., 2012). For instance, substantial genetic variation of the common tree *Corymbia calophylla* for resistance to the leaf blight pathogen *Quambalaria pitereka* was found in Australia (Ahrens et al., 2019). To investigate the genetic architecture underlying the adaptive potential of the TOU-A population in response to more southern strains of the *P. syringae* complex, we focused on the JACO-CL strain that induces both severe symptoms on TOU-A plants in comparison to accessions from its native area and variable disease responses among TOU-A accessions (Figure 2, Table S2). As a first step, we generated a *de novo* genome sequence of the JACO-CL strain.

### The JACO-CL strain belongs to the *Pseudomonas syringae* Phylogroup 7 *clade a* and displays flexible genomic traits

The genome of the JACO-CL strain was sequenced by coupling the Oxford Nanopore Technology (ONT) and the Illumina technology. The assembly is composed by two circularized sequences (one chromosome and one plasmid), resulting in 6,043,462 bp for the chromosome and 42,253 bp for the plasmid. The genome annotation resulted in 5,352 CDS, 16 rRNA, 70 tRNA and 2 tmRNA. Genome annotation and genomic analysis suggest genome plasticity and flexibility of the JACO-CL strain. First, we found two prophages (JACO-CL PP1 and JACO-CL PP3) displaying similarity with PHAGE_Vibrio_VP58.5_NC_027981 and PHAGE_Salmon_118970_sal3_NC_031940 (Supplementary Data Set 2). Secondly, we identified the 42,253 bp plasmid, thereby stressing the JACO-CL genome plasticity. KEGG orthology analysis on the plasmid open reading frames (ORFs) shows that most of the plasmidic genes encode for VirB type IV secretion system (T4SS) subunits (VirB1, VirB2, VirB4, VirB5, VirB6, VirB9, VirB10, VirB11) forming the *virB* operon (Supplementary Data Set 3). The *virB* operon was described on the Ti plasmid of *Agrobacterium tumefaciens*. Structural analysis evidenced eleven ORFs coding for potential membrane-spanning regions or signal peptides (Shirasu et al., 1990). Interestingly, the *virB* operon was also observed and extensively studied in the phytopathogenic *Xanthomonas* species. For instance, the role of the plasmid in pathogen virulence has been investigated in *virB7* knockout *X. citri* mutants and showed no effect on the development of canker symptoms in citrus (Souza et al., 2011). Similar results were obtained on *X. campestris* pv. *campestris*, with the deletion of the *virB* plasmid that did not modify phytopathogen aggressiveness on Brassica crop species (He et al., 2007). Conversely, the ability of the *VirB* T4SS in killing Gram-negative bacteria has been demonstrated in both *X. citri* (Souza et al., 2015) and *Salmonella maltophilia* (Bayer-Santos et al., 2019). Indeed, T4SS is responsible for transferring toxic effectors into the cells of bacterial competitors. Altogether, the presence of the T4SS *virB* plasmid in JACO-CL corroborates with the hypothesis that this strain might employ T4SS effectors to compete with bacterial populations inhabiting *A. thaliana* phyllosphere. This might be an additional evidence of the evolutionary history of *P. syringae* PG7 as a widely spread and common bacterium co-evolving with Arabidopsis.

Functional studies validating the role of the *virB* T4SS will help in clarifying whether or not this operon is a crucial determinant of competitive interactions between *P. syringae* and members of *A. thaliana* microbiota. Studies on the occurrence of the T4SS *virB* operon in *P. syringae* populations inhabiting *A. thaliana* will also contribute to understand the role of T4SS on the evolutionary forces driving the emergence of novel highly competitive pathogens.

### The JACO-CL strain carries a Single-Pathogenicity Island (S-PAI) T3SS and hop genes potentially related with pathogenicity in *A. thaliana*

Two PG7 genotypes with distinct virulence phenotypic traits were isolated in the past from *A. thaliana* (Araki et al., 2006; Jakob et al., 2007). In particular, PG7 natural populations carry two structurally divergent Pathogenicity Islands (PAIs), which encode for T3SS genes and are located in different chromosomal locations. The T-PAI is organized as the classical *P. syringae* tripartite T3SS with an *hrp/hrc* gene cluster, the exchangeable effector locus (*EEL*) and the conserved effector locus (CEL). On the other hand, the S-PAI is composed of the *hrp/hrc* cluster containing the *avrE* effector and its chaperone. Interestingly, PG7 isolates only possess one of the PAI as a possible result of balancing selection on the pathogen associated with the presence/absence of specific host resistance genes (Araki et al., 2006). To investigate the type of JACO-CL PAI, we first inferred a phylogeny based on the *avrE* effector located in the JACO-CL *hrp/hrc* gene cluster and reference PG7 strains carrying the S-or T-PAI island (Figure 3a). The *avrE* is a conserved effector of the *hrp/hrc* cluster present on both S-and T-PAI strains. However, *avrE* divergently evolved in the two distinct PAIs. A phylogenetic tree shows that JACO-CL carries a S-PAI configuration and this was confirmed by genome structural analysis (Figures 3a and 3b, Supplementary Data Set 4). In addition, KEGG analysis also corroborates that T3SS is not complete in JACO-CL as expected for S-PAI strains that have a reduced T3SS (Supplementary Data Set 5). More specifically, JACO-CL harbors only a *hrp/hrc* cluster containing both *avrE* and *avrF* and their relative chaperons (Figure 3b).

**Figure 3.**
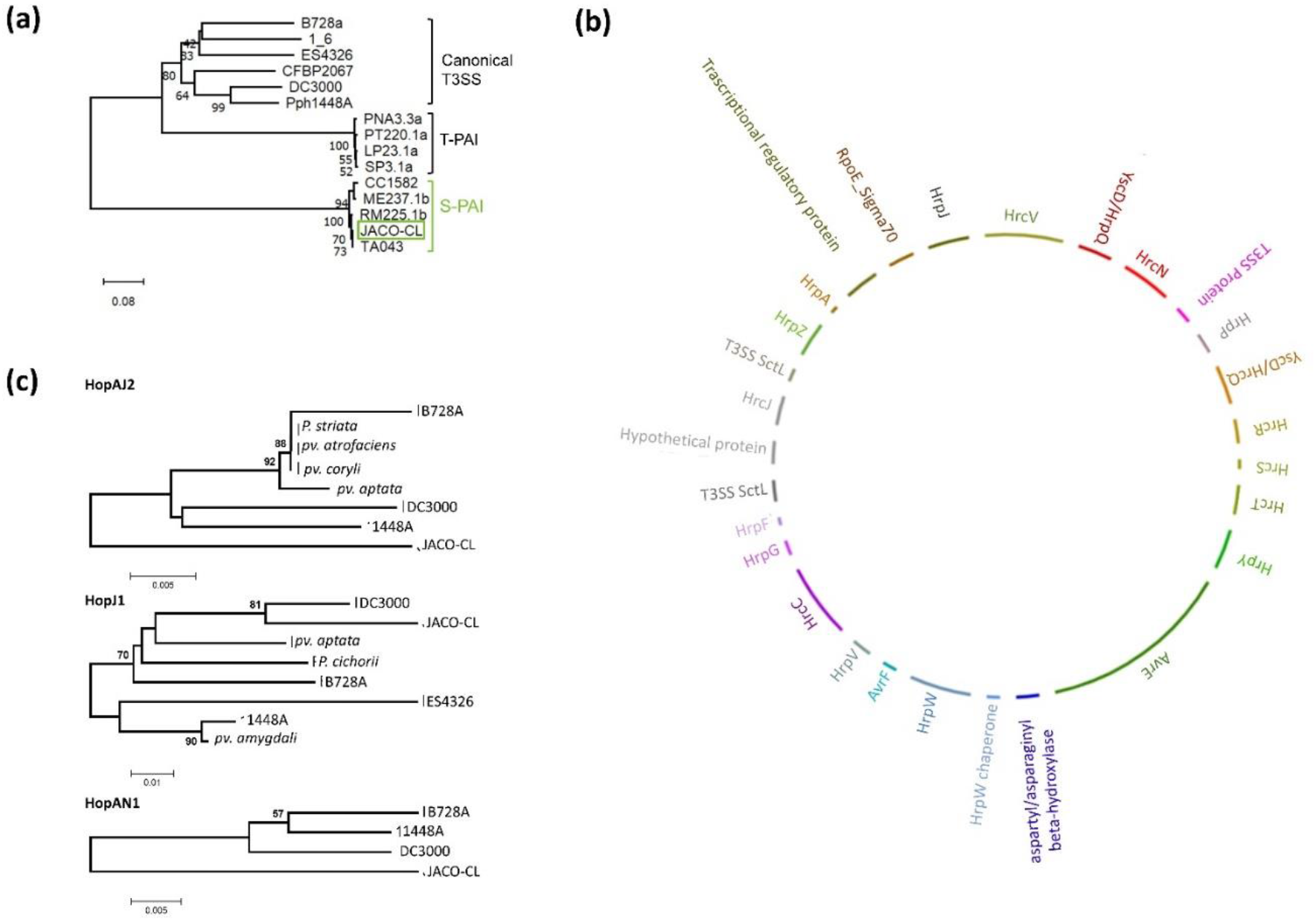
Type III Secretion System of the JACO-CL strain. **(a)** Phylogenetic tree (Neighbor joining, NJ) based on the nucleotide sequence of the *avrE* gene (5160 bp). T-PAI and S-PAI (indicated in red) *Pseudomonas viridiflava* strains were integrated in the analysis to investigate the T3SS JACO-CL affiliation. DC3000 (PG1), B728a (PG2), 1_6 (PG4), ES4326 (PG5), CFBP2067 (PG6) *avrE* nucleotide sequences were included in the analysis as reference strains carrying a canonical T3SS. The phylogenetic analysis demonstrates that the JACO-CL strain carries a S-PAI T3SS that is closely related to the TA043 strain isolated from *Primula officinalis* (Bartoli *et al*., 2014). **(b)** JACO-CL hrp/hrc Type III locus reconstructed after genome annotation. Line lengths are proportional to protein size and protein annotation was performed both based on Bacannot pipeline and manual correction. **(c)** Phylogenetic inference (Neighbor Joining, NJ) on the amino acid sequences of the predicted Hop proteins found in the JACO-CL strain. Reference protein sequences were recovered on Pseudomonas-Plant Interaction (PPI) database http://www.pseudomonas-syringae.org/pst_func_gen2.htm) and GeneBank. NJ trees show that all hops (HopAJ2, HopJ1 and HopAN1) from JACO-CL are divergent from those described previously in other *Pseudomonas syringae* pathovars.

Interestingly, through genome functional annotation, we found three complete Hop effectors (HopAJ2, HopAN1 and HopJ1) located in the genome of JACO-CL but not physically related with the PAI (Supplementary Data Set 4). To investigate the possible Horizontal Gene Transfer (HGT) of the tree Hop effectors from other *P. syringae* PGs, we compared the JACO-CL alleles with those already described in *P. syringae* and available at the Pseudomonas-Plant Interaction (PPI) http://pseudomonas-syringae.org. We also added alleles retrieved from Genebank (BLASTP > 50% of protein coverage). Phylogenetic relationships between the Hop protein sequences indicate that both HopAJ2 and HopAN1 are highly divergent from alleles previously described in other *P. syringae* PGs (Figure 3c). This result indicates that both Hop effectors were not recently acquired through HGT from existing alleles distributed in the *P. syringae* complex. On the opposite, phylogeny based on the HopJ1 protein sequences indicates that JACO-CL HopJ1 allele is closely related to the allele found in DC3000 strain (Figure 3c) (*P. syringae* pv. *tomato*, PG1 *clade a*), thereby suggesting a possible HGT acquisition.

HopAJ2_Pph1448a_ and HopAN1_Pph1448a_ alleles were tested for translocation. Pph1448a strain (*P. syringae* pv. *phaseolicola*) belongs to *P. syringae* PG3. Standard Hypersensitive Response (HR) and competitive assays were performed and none of the Pph1448a candidate fusions elicited HR, thereby proving the absence of translocation of Pph1448a alleles (Macho et al., 2009). This result indicates that HopAJ2 and HopAN1 should be considered as putative effectors. However, the differences between the JACO-CL protein sequences compared to Pph1448a (Figure 3c) suggest that HopAJ_2JACO-CL_ and HopAN1_JACO-CL_ might be still functional. Translocation assays in tobacco and Arabidopsis are necessary to validate these putative effectors in *P. syringae* strains with a non-canonical T3SS. Up to now, the role of these Hop proteins remains speculative but opens new perspectives in the validation of novel effectors. Regarding HopJ1, its translocation was observed in DC3000 but not in Pph1448a (Macho et al., 2009). However, HopJ1_DC3000_ and HopJ1_Php1448a_ show N-terminal changes sufficient to explain translocation variability. On the other hand, HopJ1_JACO-CL_ shows a high protein similarity with the HopJ1_DC3000_, thereby making this effector a good candidate for validation assays in *P. syringae* PG7. Investigation of the prevalence of these effectors in natural *A. thaliana* populations will also be interesting to understand their role in the evolution of pathogenicity in the *P. syringae* complex.

### The adaptive potential for disease resistance is mediated by a polygenic architecture

To investigate the genetic architecture underlying the adaptive potential of response to the *P. viridiflava* strain JACO-CL, we inoculated 192 whole-genome sequenced A. thaliana accessions of the TOU-A population. Quantitative variation in disease resistance to the JACO-CL strain was observed between the 192 TOU-A accessions (Supplementary Data Set 6). Similar quantitative variation among TOU-A accessions was observed in response to either the bacterial pathogens *Xanthomonas campestris* pv *campestris* (*Xcc*) and *Ralstonia solanacearum* (Debieu et al., 2016; Demirjian et al., 2022; Desaint et al., 2021; Huard-Chauveau et al., 2013) or natural infections in the native habitats of the TOU-A population (Roux & Frachon 2022).

Highly significant genetic variation was observed at 1 dai (accession effect: λ_LR_ = 38.0, *P* = 7.07E-10), 2 dai (accession effect: λ_LR_ = 70.8, *P* < 1.0E-16) and 3 dai (accession effect: λ_LR_ = 64.0, *P* = 1.22E-15) (Figure 4), with broad-senses heritability values of 0.48, 0.58 and 0.57 at 1 dai, 2 dai and 3 dai, respectively. These results suggest that a non-negligible fraction of the adaptive potential of response to the JACO-CL strain is mediated by host genetics. Similar results were obtained for the pathosystems *Fraxinus excelsior* - *Hymenoschyphus* (Kjær et al., 2012), *Pinus pinaster* – *Fusarium circinatum* (Elvira-Recuenco et al., 2014) and *C. calophylla* – *Q. pitekera* (Ahrens et al., 2019). However, narrow-sense heritability values were smaller for *F. excelsior* (∼0.36) and *C. calyphylla* (∼0.14) than for *Pinus pinaster* (∼0.53), thereby highlighting variation among species in the adaptive genetic potential to face novel or emerging pathogens.

**Figure 4.**
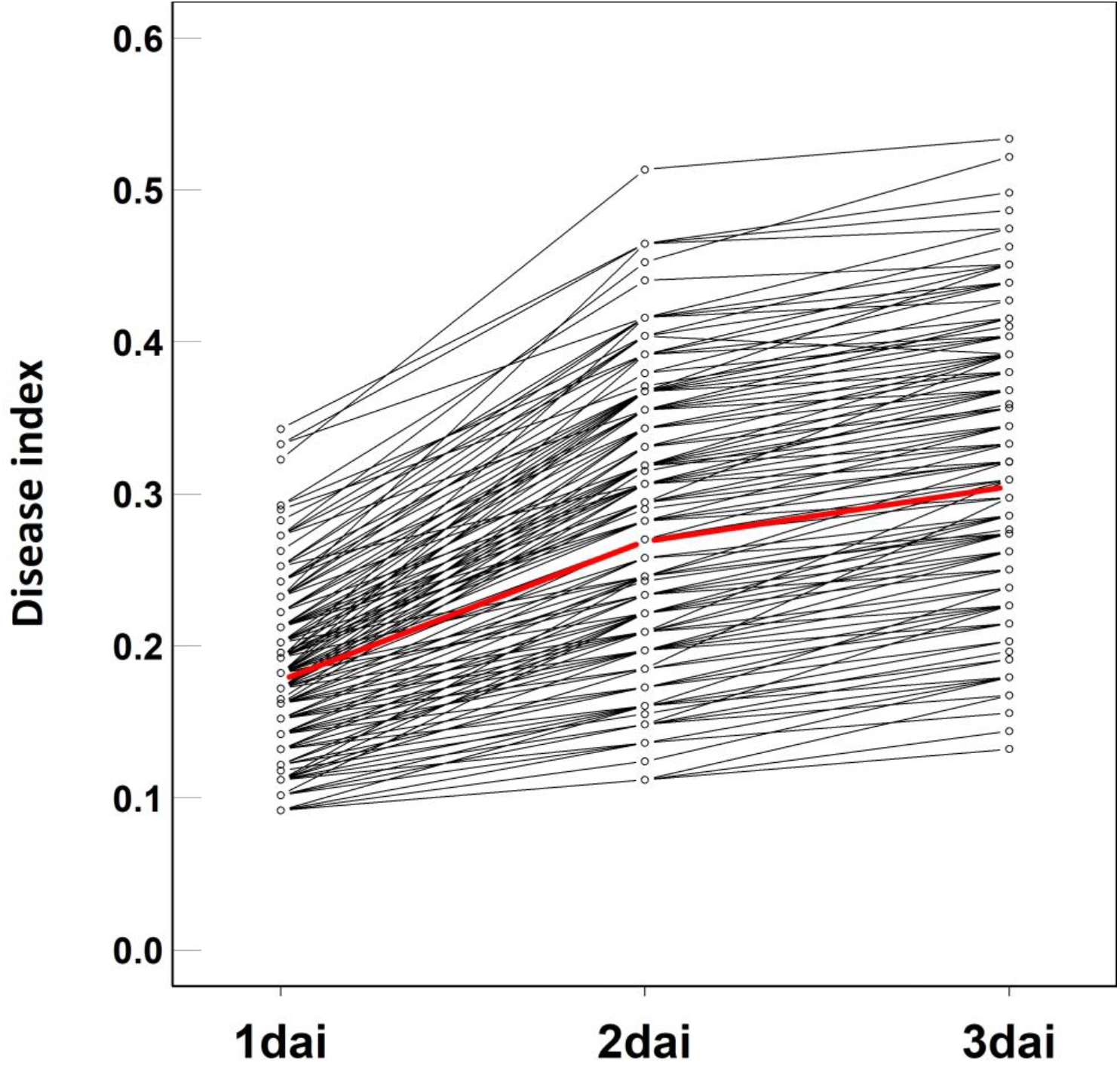
Genetic variation of plant response in the TOU-A local mapping population to the JACO-CL strain at 1dai, 2dai and 3dai. The dots correspond to the genotypic values of the 192 accessions. Each black line corresponds to the plant response to one of the 192 accessions. The red line represents the mean of disease index among the 192 accessions.

To fine map QTLs associated with quantitative disease resistance (QDR) to the JACO-CL strain, we took advantage of the Illumina genome sequencing of the 192 TOU-A accessions phenotyped in this study, which revealed a set of 1,902,592 Single Nuclear Polymorphisms (SNPs) and a linkage disequilibrium (LD) decay to r^2^ = 0.5 within an average of 18 bp (Frachon et al., 2017). We combined a mixed-model approach correcting for the effects of population structure with a local score approach, which allows delimiting QTL regions by accumulating single marker *p*-values obtained from the mixed-model while controlling the issue of multiple hypothesis tests (Bonhomme et al., 2019). This combined GW-LS approach was previously demonstrated to be efficient in the TOU-A population, with the fine mapping (in particular, the detection of QTLs with small effects) and functional validation of one QTL of QDR to *Xcc* (Huard-Chauveau et al., 2013) and one QTL of QDR to *R. solanacearum* (Aoun et al., 2020). QDR to the JACO-CL strain was mediated by a polygenic architecture (Figure 5), with the detection of 16 QTLs, 14 QTLs and 22 QTLs at 1 dai, 2 dai and 3 dai, respectively (Supplementary Data Set 7). All these QTLs overlap with 87 unique candidate genes (Supplementary Data Set 7). Almost half of the QTLs were common between at least two days of scoring, whereas the other QTLs were specific to one day of scoring (Figure 6), thereby suggesting that the adaptive potential of response to the JACO-CL strain is mediated by a mix of constitutive and dynamic QTLs.

**Figure 5.**
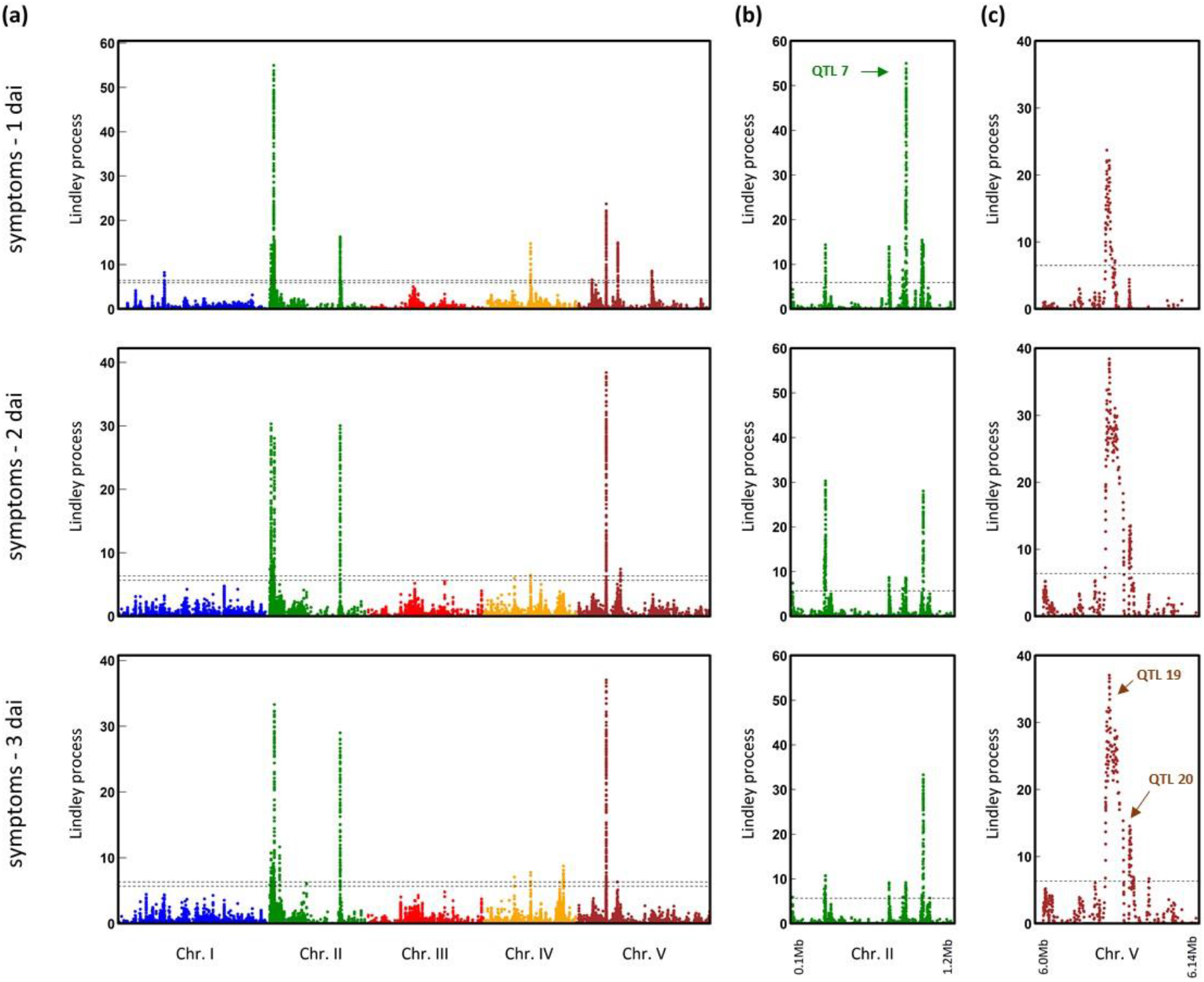
The genetics of quantitative disease resistance to the JACO-CL strain in the TOU-A population. **(a)** Manhattan plots comparing the GWA mapping results at 1 dai, 2 dai and 3 dai. The x-axis indicates the physical position of the 978,408 SNPs on the five chromosomes. The y-axis indicates the Lindley process scores estimated from -log_10_ *p*-values from the mixed model implemented in the software EMMAX using SNPs with a minor allele relative frequency (MARF) > 7%. **(b)** Zoom spanning a genomic region at the beginning of chromosome II from 0.1Mb to 1.2Mb. **(c)** Zoom spanning a genomic region at the beginning of chromosome V from 6Mb to 6.14Mb. The dashed line indicates the maximum of the five chromosome-wide significance thresholds. Arrows indicate the three QTLs containing candidate genes of particular interest.

**Figure 6.**
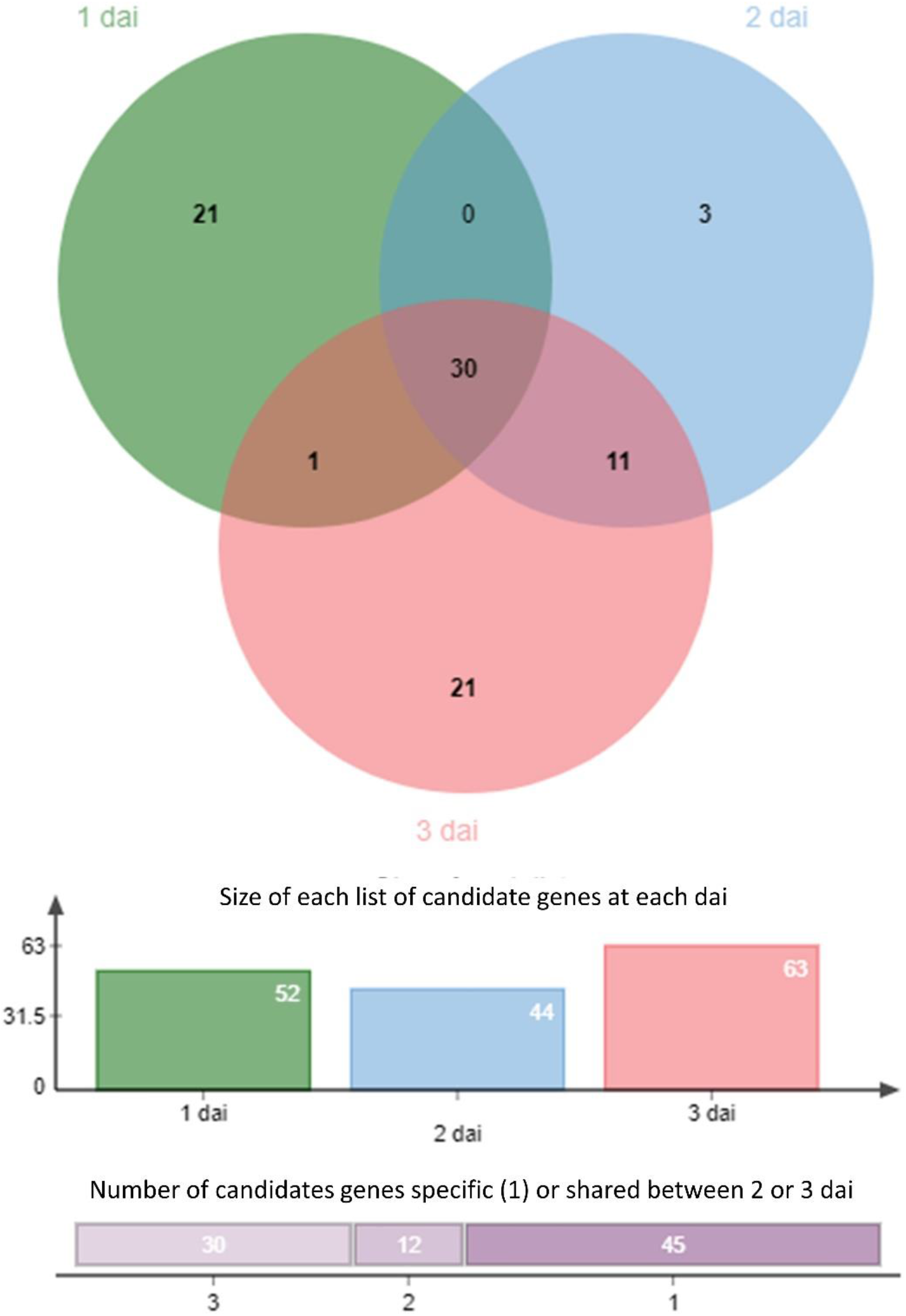
Venn diagram illustrating the number of common and specific candidate genes between the three days of scoring. Colored bars indicate the number of candidate genes at each dai. The vertical stacked bar plot indicates the number of candidate genes specific to one dai (right) or shared between two or three dai.

Two constitutive QTLs overlap with candidate genes of particular interest. The first QTL correspond to 95 top SNPs (at 1 dai) located on chromosome 2 (QTL7, Figure 5b) in the vicinity or centered on the gene *RIBONUCLEASE 1* (*RNS1, AT2G02990*) (Supplementary Data Set 7), which encodes a member of the ribonuclease T2 family that is involved in wound-induced signaling independent of jasmonic acid (LeBrasseur et al., 2002). The second QTL corresponds to 44 top SNPs (at 3 dai) located on chromosome 5 (QTL19, Figure 5c) in the vicinity or centered in the gene *AT5G18310*, which encodes for a ubiquitin hydrolase. Interestingly, this protein is the target of the effector AvrRpt2 of *P. syringae* (Chisholm et al., 2005). A 4.04-fold variation of the expression level of *AT5G18310* in leaves was observed between 665 worldwide accessions of *A. thaliana* (Supplementary Data Set 8) (Kawakatsu et al., 2016). GWA mapping revealed that the natural variation in the expression of *AT5G18310* is controlled at the worldwide scale by one QTL located in the 5’ region of *AT5G18310*, with top SNPs that were common with the top SNPs identified in response to the JACO-CL strain (Figure S1). This suggests that modification of cis-regulation of *AT5G18310* mediates genetic variation of the response to the JACO-CL strain in the TOU-A population. Estimating the expression level of *AT5G18310* in TOU-A accessions contrasted in their response to the JACO-CL strain could support this hypothesis. In addition, *AT5G18130* is located nearby a dynamic QTL (QTL20, Figure 5c) encompassing two genes (*AT5G18350* and *AT5g18360*) encoding for Toll/Interleukin-1 receptor nucleotide-binding site leucine-rich repeat (TIR-NBS-LRR) proteins. Based on the Illumina sequencing coverage depth of the QTL 20, no presence/absence (P/A) polymorphism was detected for *AT5G18360*. However, we detected a P/A polymorphism for *AT5G18350* (present in 134 accessions and absent in 58 accessions), which is line with the P/A polymorphism for *AT5G18350* identified on a set of 33 worldwide accessions by PCR amplification (Shen et al., 2006). Given the function of the candidate genes underlying these two closely located QTLs on chromosome 5, it is tempting to propose a scenario similar to the one observed between the effector AvrRpt2, the guard protein RIN4 and the *R* gene RPS2, where degradation of RIN4 proteins by the proteolytic activity of AvrRpt2 triggers defense responses *via* RPS2 (Chisholm et al., 2005). In our study, AT5G18310 proteins might be degraded by the proteolytic activity of one the three complete Hop effectors (HopAJ2, HopAN1 and HopJ1), which in turn would induce defense responses *via* the *R* genes AT5G18350/AT5G18360. Further experiments testing protein-protein interactions are needed to validate this scenario.

Because genetic variation was found for disease resistance to the *P. syringae sensus stricto* strain RAYR-BL (Table S2), it would be interesting to test whether genetic solutions underlying the adaptive potential of response of the TOU-A population are similar between strains of *P. syringae sensu stricto* and *P. viriflava* originating from more southern geographic regions. It would definitely help to better understand the genetic and molecular basis of the adaptive potential in ecologically relevant *A. thaliana* – *P. syringae* pathosystems.

## EXPERIMENTAL PROCEDURES

### Bacterial strains and phylogroup affiliation

The eight strains considered in this study were isolated from eight natural populations of *A. thaliana* located south-west of France (Figure 1A) (Bartoli et al., 2018). Briefly, in spring 2015, two or three plants of each natural population of *A. thaliana* were transferred to the laboratory and leaves were aseptically homogenized and serial dilutions were plated on Trypticase Soy Agar (TSA) medium supplemented with 100 mg/L of cycloheximide. To detect *P. syringae*-like strains, isolates were screened by PCR with a *Psy* marker (Guilbaud et al., 2016). All the strains positive for the *Psy* marker were amplified with the primer set *shcA* described in (Bartoli et al., 2014), which allows discriminating among PG 7 *P. syringae* haplotypes.

Phylogroup/clade affiliation of the eight strains was defined based on the *P. syringae* species complex nomenclature (Berge et al., 2014). To do so, the sequences of the housekeeping gene *citrate synthase* (*cts*) obtained in (Bartoli et al., 2018) and the reference *cts* nucleotide sequences of *P. syringae* obtained from (Berge et al., 2014) were aligned with DAMBE version 5.1.1 (Xia, 2013). MEGA 7 was used to infer the phylogeny by using a maximum likelihood model (Kumar et al., 2016).

### Genome sequencing of the JACO-CL strain

High Molecular Weight DNA of the JACO-CL strain was extracted from an overnight culture (TSA medium, 28°C) following the protocol of Mayjonade et al. (dx.doi.org/10.17504/protocols.io.r9id94e). An Oxford Nanopore Technology (ONT) library was prepared using the EXP-NBD103 and SQK-LSK108 kits according to the manufacturer’s instructions and starting with 3 μg of 20 kb sheared DNA (Megaruptor, Diagenode) as input. This library was pooled with 11 other samples and 100 fmol of this pool was loaded and sequenced on one R9.4.1 flowcell for 72 h. The same DNA sample was used to prepare an Illumina library using the TruSeq Nano DNA HT Library Prep Kit (Illumina). This library was sequenced with 43 other samples on a Hiseq3000 instrument (2×150 bp paired-end reads) (Illumina) at the GeT-PlaGe INRAE Sequencing facility (Toulouse, France).

Nanopore raw signals were filtered with Guppy 2.1.3. After filtering, we obtained 52, 026 filtered reads (min = 215, max = 49, 692) corresponding to 654, 999, 490 bp. The filtered reads were assembled with Canu 1.7 (Koren et al., 2017) with default parameters. Circularization of the assembled genome was performed with a home-made pipeline running a NCBI-blastn by filtering with a minimal length of the High-scoring Segment Pair (HSP) – a local alignment without gaps – of 10, 000 nt and an overhand of 5, 000 nt. Illumina sequencing yielded 1, 277, 516 paired-end reads corresponding to 381, 654, 800 bp. The Nanopore assembly was error corrected by mapping the Illumina reads with BWA (version 0.7.17) and then performing two rounds of polishing with Pilon (version 1.22). Finally, we obtained two circular JACO-CL sequences, one corresponding to a plasmid (42, 253 bp) and one corresponding to the genomic DNA of the strain (6, 043, 462 bp). Genomic DNA and plasmid were deposited in GenBank under Accession Numbers CP097286 and CP097286 respectively (related Bio Sample SAMN28171611and Bio Project PRJNA836746).

### Genomic characterization of the JACO-CL strain

The circular JACO-CL genome was annotated and analyzed with the Bacannot framework https://bacannot.readthedocs.io/en/latest/#index--page-root using Prokka version 1.14.5 for annotation (Seemann, 2014). KEGG KO annotation was performed with KofamScan version 1.3.0 (Aramaki et al., 2020). KEGG analysis was separately run on genomic and plasmid DNA sequences and plasmid protein annotation was checked by BLASTP by searching for protein sequences showing homology with known plasmids.

PhiSpy and PHASTER were used for phage finding and annotation (Akhter et al., 2012; Zhou et al., 2011). T3SS effector proteins identification was determined through BLASTP search of the effector proteins reported at the Pseudomonas-Plant Interaction (PPI) http://pseudomonas-syringae.org database with an e-value of 1e-6. We enriched this existing effector database with T3SS effector proteins reported previously in the *P. syringae* PG7 (Bartoli et al., 2014). Only complete effectors (alignment length ≥ 25%) were kept in the analysis. In addition, putative effector proteins and/or proteins found in the CEL were manually blasted in GenBank for further annotation. Phylogenetic relationships for the three complete Hop proteins (HopAJ2, HopAN1 and HopJ1) found in the JACO-CL genome were estimated by comparing these Hops with alleles retrieved at the PPI database and on GenBank through BLASTP of the JACO-CL Hop sequences. Amino acid sequences of each Hop were individually aligned with MAFFT v7.271 (Katoh and Standley, 2013)and a Maximum Likelihood (ML) tree was inferred with MEGA7 (Kumar et al., 2016).

### Genotype x Genotype interactions

To estimate the genetic potential of the local mapping population TOU-A to face strains of the *P. syringae* complex, seven TOU-A accessions (A1-9, A1-19, A1-137, A6-18, A6-47, A6-55 and A6-105) and the reference accession Col-0 were inoculated with each of the eight strains (BAZI-AL, JACO-CL, NAZA-AL, NAUV-BL, RADE-A4, RAYR-BL, SIMO-AL and SAUB-A). The reference accession Col-0 was also inoculated with the same set of eight strains. A growth chamber experiment with 192 plants was set up at the Toulouse Plant Microbe Phenotyping Platform (TPMP), with three plants inoculated for each ‘accession*strain’ combination. After a 4-day stratification treatment, plants were grown in Jiffy pots at 22 °C under 90% humidity and artificial light to provide a 9-hr photoperiod (Huard-Chauveau et al., 2013). Bacterial inoculation was conducted on 28-day-old plants using a blunt-ended syringe (Terumo® SYRINGE 1mL, SS+01T1). Four leaves per plant were entirely infiltrated with a 5.10^7^ CFUmL-1 bacterial solution. Disease symptoms were visually scored at 1, 2, 3 and 4 dai as described in (Roux et al., 2010). Each infected leaf received a score from 0 to 1, with 0 corresponding to no symptoms whereas 0.5 and 1 correspond to medium and severe symptoms, respectively. These scores are determined by the presence of visible chlorosis, visible necrosis, leaf mosaic or water-soaked lesions and cell death related symptoms surrounding infection sites. We averaged the scores of the four inoculated leaves per plant.

For each of the six strains of the *P. syringae* complex for which symptoms were observed on plants (Supplementary Table S1), we explored genetic variation among accessions at each dai, using the following model:

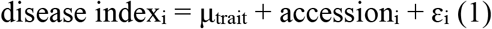

where ‘μ’ is the overall phenotypic mean; ‘accession’ accounts for differences among the eight *A. thaliana* accessions; ‘ε’ is the residual term. Inference was performed using ReML estimation, using the PROC MIXED procedure in SAS v.9.4 (SAS Institute Inc., Cary, North Carolina, USA). The ‘accession’ factor was treated as a fixed effect. Because *A. thaliana* is a highly selfing species (Platt et al., 2010), Least-squares means (LS-means) obtained for each accession from model (1) correspond to the genotypic values of accessions.

To explore genetic variation among the eight strains, we used the following model at each dai:

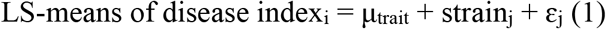

where ‘μ’ is the overall mean of LS-mean values; ‘strain’ accounts for differences among the six strains of the *P. syringae* complex for which symptoms were observed on plants; ‘ε’ is the residual term. Inference was performed using ReML estimation, using the PROC MIXED procedure in SAS v.9.4 (SAS Institute Inc., Cary, North Carolina, USA). The ‘strain’ factor was treated as a fixed effect.

### Genome-Wide Association mapping of response to the JACO-CL strain

#### Experimental design and growth conditions

To investigate the genetic architecture underlying the adaptive potential of response to the JACO-CL strain, we inoculated 192 whole-genome sequenced accessions of the TOU-A population. A growth chamber experiment with 1,152 plants was set up at the Toulouse Plant Microbe Phenotyping Platform (TPMP) using a randomized complete block design (RCBD) with six experimental blocks. Each block was represented by 192 plants corresponding to the 192 TOU-A accessions. Growth chamber conditions, bacterial inoculation and disease scoring at 1 dai, 2 dai and 3 dai, were similar to the experiment conducted on the eight accessions, as described above.

#### Statistical analysis

At each dai, we explored genetic variation among the 192 TOU-A accessions using the following model:

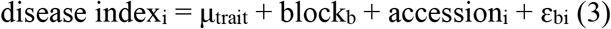

where ‘μ’ is the overall phenotypic mean; ‘block’ accounts for differences in micro-environment among the six experimental blocks; ‘accession’ accounts for differences among the 192 TOU-A *A. thaliana* accessions; ‘ε’ is the residual term. Inference was performed using ReML estimation, using the PROC MIXED procedure in SAS v.9.4 (SAS Institute Inc., Cary, North Carolina, USA). While the ‘block’ factor was treated as a fixed effect, the ‘accession’ effect was treated as a random effect. Significance of the random effect was determined by likelihood ratio tests of the model with and without this effect. Best Linear Unbiased Estimates (BLUEs) of disease index were obtained for each accession from model (3). Broad-sense heritability (*H*^2^) of disease index was estimated from variance component estimates for the ‘block’ and ‘acc’ terms (PROC VARCOMP procedure in SAS v.9.4)

#### GWA mapping combined with a local score approach

The effects of population structure on phenotype-genotype associations were demonstrated to be limited in the TOU-A population (Baron et al., 2015; Brachi et al., 2013)]. We nonetheless run GWA mapping using a mixed model implemented in the software EMMAX (Efficient Mixed-Model Association eXpedited) (Kang et al., 2010). To control for the effect of population structure, we included as a covariate an identity-by-state kinship matrix K. This kinship matrix was based on 1,902,592 SNPs identified among the 192 accessions of the TOU-A population (Frachon et al., 2017). Because rare alleles may increase the rate of false positives (Atwell et al., 2010; Bergelson and Roux, 2010; Brachi et al., 2010), we only considered SNPs with a minor allele relative frequency (MARF) > 7%, resulting in 978,408 SNPs. Above this MARF threshold, *p*-value distributions obtained from the EMMAX mixed model are not dependent on MARF values in the TOU-A population (Frachon et al., 2017).

In order to better characterize the genetic architecture associated with natural genetic variation in response to JACO-CL, a local score approach was applied on the set of *p*-values provided by EMMAX. This local score approach increases the power of detecting QTLs with small effect and narrows the size of QTL genomic regions (Bonhomme et al., 2019; Fariello et al., 2017). In this study, we used a tuning parameter ξ of 2 expressed in –log_10_ scale. Significant phenotype-SNP associations were identified by estimating a chromosome-wide significance threshold for each chromosome (Bonhomme et al., 2019). Based on a custom script (Libourel et al., 2021), we retrieved all candidate genes underlying QTLs by selecting all genes inside the QTL regions as well as the first gene upstream and the first gene downstream of these QTL regions (Dataset 7). The TAIR 10 database (https://www.arabidopsis.org/) was used as our reference. The number of candidate genes that were either specific to a single dai or common between several dai were illustrated by a Venn diagram using the package *jvenn* (Bardou et al., 2014).

### GW-LS on the level of expression of the candidate gene AT5G18310

Data of the expression level of *AT5G18310* were retrieved from (Kawakatsu et al., 2016) for 665 whole-genome sequenced worldwide accessions that grew in similar growth conditions. Based on a set of 11,769,920 SNPs (1001 Genomes Consortium. 2016), we applied the same GW-LS approach than for investigating the genetic architecture of disease index in the TOU-A population. We considered SNPs with a minor allele relative frequency (MARF) > 10%, resulting in 1,703,894 SNPs.

## ACKNOWLEDGMENTS

This work was funded by the Région Midi-Pyrénées (CLIMARES project) and the LabEx TULIP (ANR-10-LABX-41). This work was performed in collaboration with the GeT core facility, Toulouse, France (http://get.genotoul.fr), and was supported by France Génomique National infrastructure, funded as part of “Investissement d’avenir” program managed by Agence Nationale pour la Recherche (contract ANR-10-INBS-09). Part of this work was carried out on the Toulouse Plant–Microbe Phenotyping facility (https://www6.toulouse.inrae.fr/tpmp/), which is part of the Laboratoire des Interactions Plantes-Microbes-Environnement (LIPME) - UMR INRA441/CNRS2594. We are grateful to Jérôme Gouzy for having conducted the assembly of the JACO-CL strain. We are also grateful to Fabrice Devoilles for his assistance during growth chamber experiments.

## DATA AVAILABILITY STATEMENT

Raw phenotypic data of disease index are available in Data Sets 1 and 6. Genomic DNA and plasmid were deposited in GenBank under Accession Numbers CP097286 and CP097286 respectively (related Bio Sample SAMN28171611and Bio Project PRJNA836746).

## SUPPLEMENTARY INFORMATION

**Table S1**. Mean ‘strain’ effect with and without considering the reference accession Col-0 at 1dai, 2 dai, 3 dai and 4 dai.

**Table S2**. Genetic variation among accessions for each strain, with and without considering the reference accession Col-0 at 1dai, 2 dai, 3 dai and 4 dai.

**Figure S1**. GWA mapping results of the expression level of the candidate gene *AT5G18310*.

## SUPPLEMENTARY DATA SETS

**Data Set 1**. Raw phenotypic data of disease index of seven TOU-A accessions and the reference Col-0 in response to four strains of *P. syringae* and four strains of *P. viridiflava*.

**Data Set 2**. Prophages found in the genome of the JACO-CL strain.

**Data Set 3**. pDR208 plasmid annotation based on the KEGG metabolic pathways.

**Data Set 4**. Type III Secretion System (T3SS) proteins predicted on the annotated genome of the JACO-CL strain.

**Data Set 5**. Heatmap on the KEGG metabolic pathways estimated for the JACO-CL strain.

**Data Set 6**. Raw phenotypic data of disease index scored on 192 TOU-A accessions in response to the JACO-CL strain.

**Data Set 7**. List of QTLs and associated SNPs detected for disease index at each dai.

**Data Set 8**. Normalized RNA-seq read counts for the candidate gene *AT5G18310* for 665 worldwide accessions.

