## Supplementary Information for "Unraveling the genetic architecture of the adaptive potential of *Arabidopsis thaliana* to face the bacterial pathogen *Pseudomonas syringae* in the context of global change"

### Unraveling the genetic architecture of quantitative disease resistance in an ecologically relevant *Arabidopsis thaliana* - *Pseudomonas syringae* pathosystem in the context of global change

Claudia Bartoli[1], Mylène Rigal[2], Baptiste Mayjonade[2] and Fabrice Roux[2]*

**Affiliations**

[1] Institute for Genetics, Environment and Plant Protection (IGEPP), INRAE, Institut Agro AgroCampus Ouest, Université de Rennes 1, Le Rheu, France

[2] Laboratoire des Interactions Plantes-Microbes-Environnement, Institut National de Recherche pour l'Agriculture, l'Alimentation et l'Environnement, CNRS, Université de Toulouse, Castanet-Tolosan, France

**Table S1. Mean 'strain' effect with and without considering the reference accession Col-0 at 1dai, 2 dai, 3 dai and 4 dai.**

| | Strain effect | | Strain effect without Col-0 | |
|---|---|---|---|---|
| dai | *F* | *P* | *F* | *P* |
| 1 | 12.8 | 1.40E-07 | 13.74 | 1.58E-07 |
| 2 | 18.7 | 9.67E-10 | 22.76 | 3.09E-10 |
| 3 | 21.16 | 1.65E-10 | 24.72 | 1.01E-10 |
| 4 | 19.35 | 5.98E-10 | 22.81 | 2.99E-10 |

**Table S2. Genetic variation among accessions for each strain, with and without considering the reference accession Col-0 at 1dai, 2 dai, 3 dai and 4 dai.**

| | | Accession effect | | Accession effect without Col | |
|---|---|---|---|---|---|
| Strain | dai | *F* | *P* | *F* | *P* |
| NAZA-AL | 1 | 0.91 | 0.5204 | 0.90 | 0.5216 |
| | 2 | 0.91 | 0.5204 | 0.90 | 0.5216 |
| | 3 | 0.91 | 0.5204 | 0.90 | 0.5216 |
| | 4 | 0.86 | 0.5588 | 0.83 | 0.5638 |
| RAYR-BL | 1 | 1.33 | 0.2990 | 0.83 | 0.5651 |
| | 2 | 2.61 | 0.0528 | 0.90 | 0.5231 |
| | 3 | 3.37 | **0.0209** | 2.65 | **0.0041** |
| | 4 | 3.79 | **0.0129** | 1.20 | 0.3603 |
| JACO-CL | 1 | 1.31 | 0.3056 | 1.69 | 0.1951 |
| | 2 | 2.94 | **0.0351** | 2.87 | **0.0488** |
| | 3 | 2.95 | **0.0346** | 2.93 | **0.0456** |
| | 4 | 2.94 | **0.0350** | 2.65 | **0.0041** |
| NAUVE-BL | 1 | 0.94 | 0.5048 | 1.04 | 0.4433 |
| | 2 | 1.25 | 0.3318 | 1.39 | 0.2848 |
| | 3 | 1.32 | 0.3019 | 1.45 | 0.2657 |
| | 4 | 0.80 | 0.6003 | 0.84 | 0.5606 |
| RADE-AL | 1 | 0.85 | 0.5624 | 1.19 | 0.3671 |
| | 2 | 1.03 | 0.4459 | 1.88 | 0.1550 |
| | 3 | 0.56 | 0.7743 | 0.77 | 0.6092 |
| | 4 | 0.62 | 0.7309 | 0.92 | 0.5087 |
| SAUB-AL | 1 | 1.27 | 0.3251 | 1.72 | 0.1890 |
| | 2 | 0.81 | 0.5885 | 1.10 | 0.4120 |
| | 3 | 0.34 | 0.9259 | 0.37 | 0.8853 |
| | 4 | 0.74 | 0.6316 | 0.77 | 0.6052 |

**Figure S1. GWA mapping results of the expression level of the candidate gene *AT5G18310*.** Zoom spanning a genomic region at the beginning of chromosome V from 6.04Mb to 6.0.8Mb. The dashed line indicates the chromosome-wide significance threshold. The solid line and the square illustrate the physical positions of the 5’ region and genic region of the candidate gene *AT5G18310*, respectively.

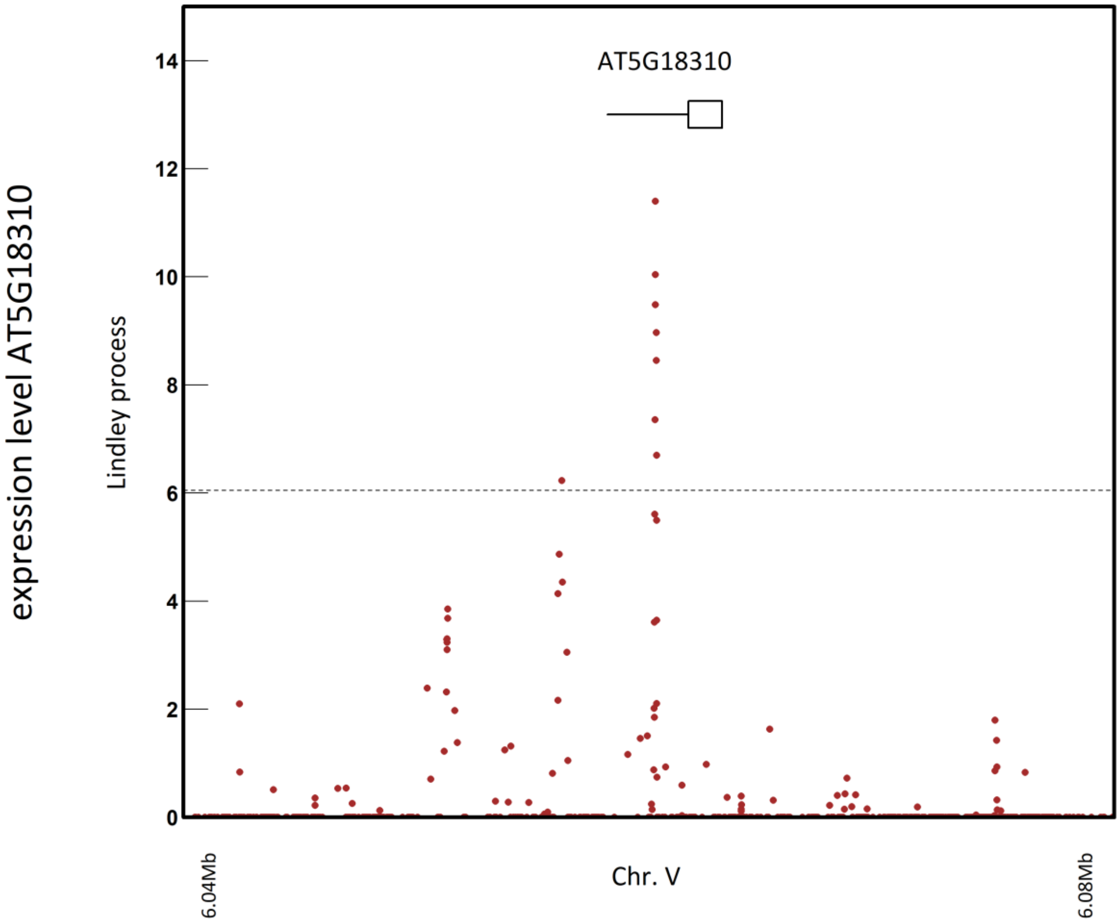
